# Rapid resistance profiling of SARS-CoV-2 protease inhibitors

**DOI:** 10.1101/2023.02.25.530000

**Authors:** Seyed Arad Moghadasi, Rayhan G. Biswas, Daniel A. Harki, Reuben S. Harris

## Abstract

Resistance to nirmatrelvir (Paxlovid) has been shown by multiple groups and may already exist in clinical SARS-CoV-2 isolates. Here a panel of SARS-CoV-2 main protease (M^pro^) variants and a robust cell-based assay are used to compare the resistance profiles of nirmatrelvir, ensitrelvir, and FB2001. The results reveal distinct resistance mechanisms (“fingerprints”) and indicate that these next-generation drugs have the potential to be effective against nirmatrelvir-resistant variants and *vice versa*.

---

Antiviral drugs are necessary to combat SARS-CoV-2/COVID-19, particularly with waning interest in the repeated vaccination boosts necessary to keep-up with virus evolution. The main protease (M^pro^) of SARS-CoV-2 is essential for virus replication and, accordingly, a proven therapeutic target as evidenced by Paxlovid (active component: nirmatrelvir; **Figure 1A**). However, as for drugs developed to treat other viruses^1^ and for first-generation SARS-CoV-2 vaccines, there is a high probability that variants will emerge that resist nirmatrelvir. Indeed, a flurry of recent studies has described a variety of candidate nirmatrelvir-resistance mutations^2–9^. Thus, considerable urgency exists to develop next-generation M^pro^ inhibitors with different resistance mechanisms and, in parallel, robust systems to rapidly assess the potential impact of candidate resistance mutations.

**Figure 1.**
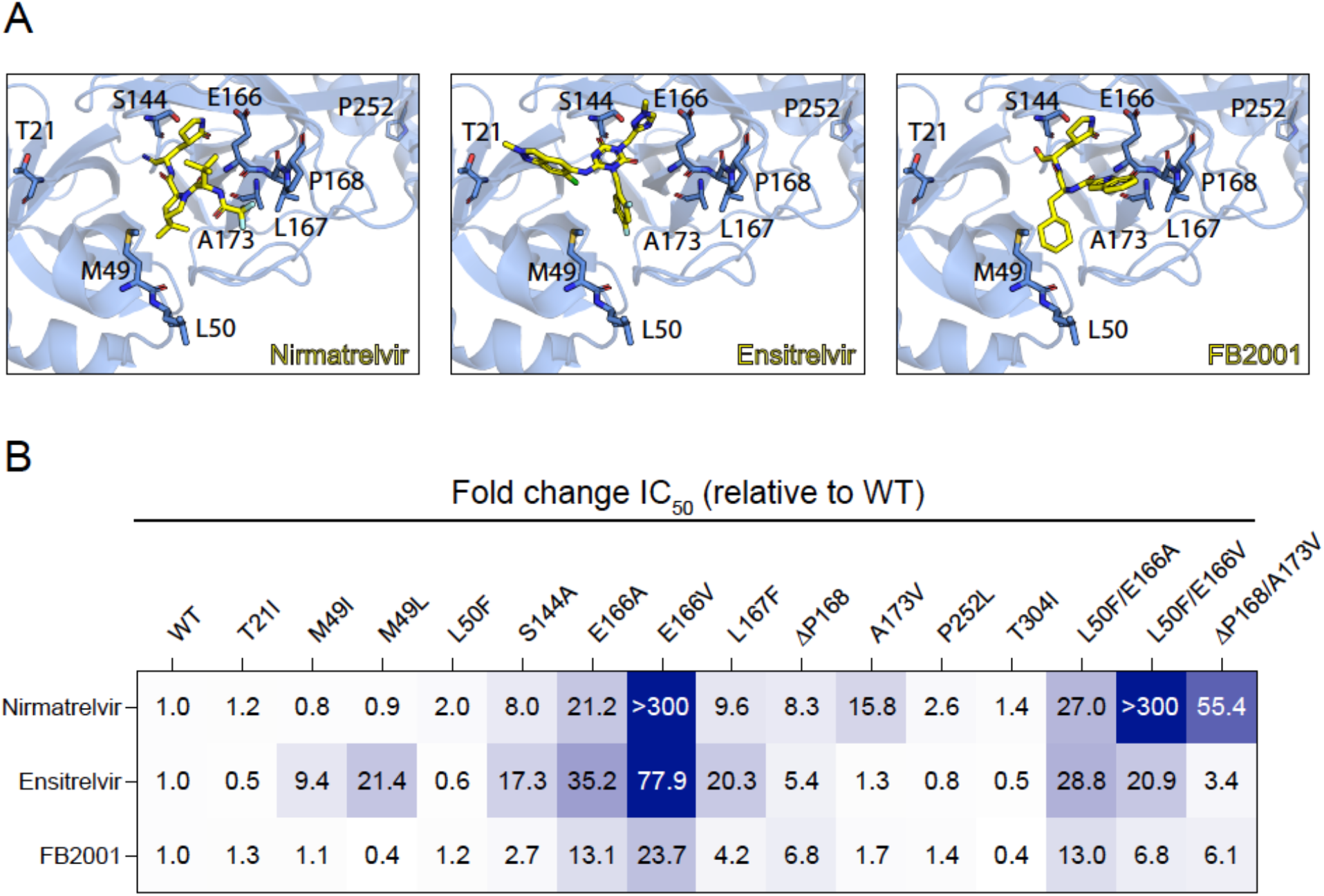
Resistance profiles of nirmatrelvir, ensitrelvir, and FB2001. (**A**) Co-crystal structures of SARS-CoV-2 M^pro^ in complex with nirmatrelvir (PDB:7SI9), ensitrelvir (PDB:7VU6), or FB2001 (PDB:6LZE). Labeled residues are interrogated in panel B. (**B**) Fold-change in IC_50_ relative to WT for the indicated mutants using the live cell gain-of-signal assay in 293T cells.

Ensitrelvir (Xocova) and FB2001 are being evaluated in clinical trials, and the former drug also recently received EUA in Japan^10,11^ (**Figure 1A**). We recently developed a gain-of-signal system for facile quantification of M^pro^ inhibition^12^, and subsequently used it together with an evolution- and structure-guided approach to characterize candidate nirmatrelvir- and ensitrelvir-resistance mutations^2^. Here, an expanded panel of M^pro^ single and double mutants based on recent studies by our group and others^2–9^ is leveraged to determine resistance profiles of these two drugs, as well as FB2001, a potential next-generation therapy (heatmap of results in **Figure 1B**; quantification summary in **Table 1**; representative dose responses in **Figure S1**).

**Table 1.**
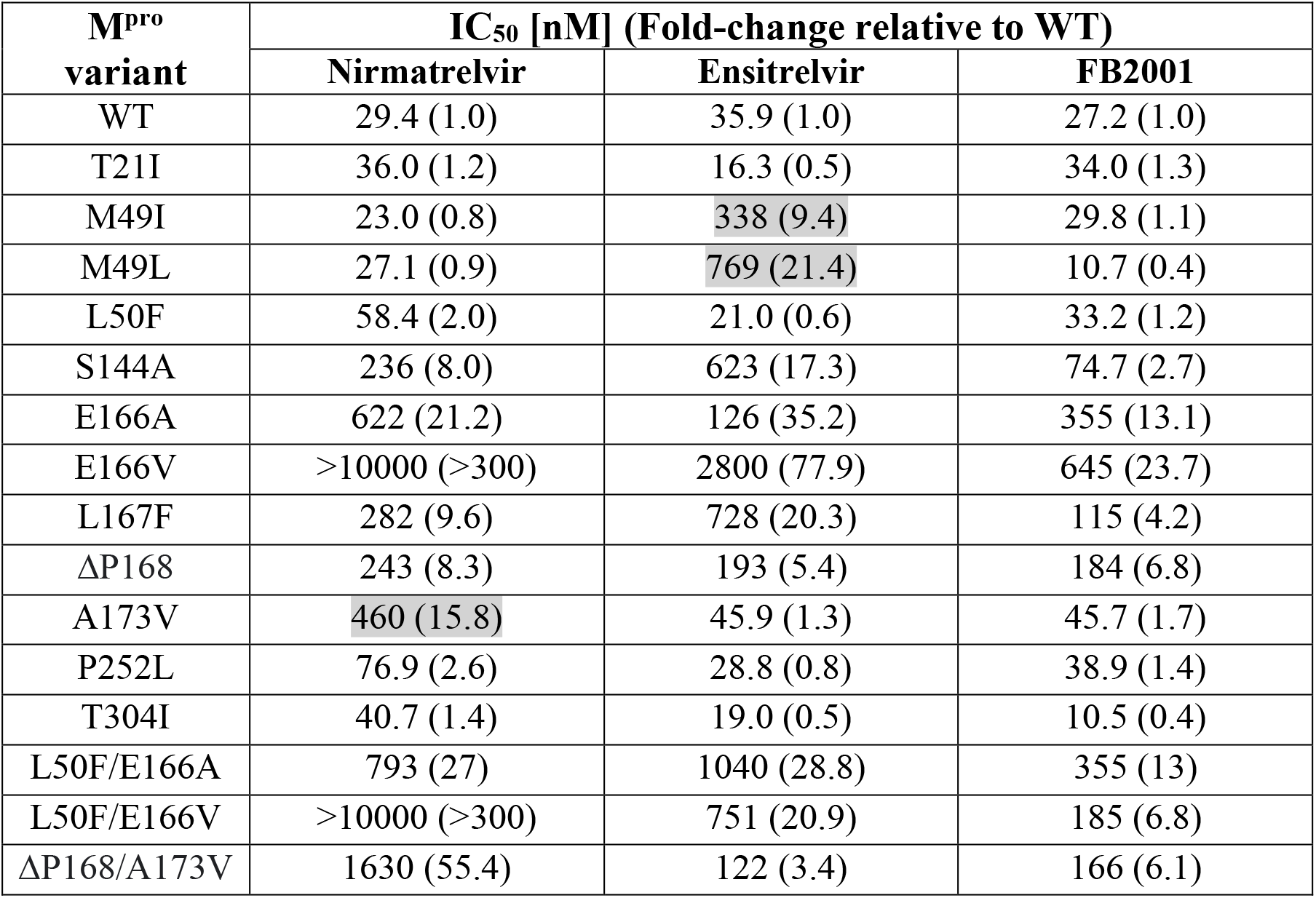
IC_50_ values of nirmatrelvir, ensitrelvir, and FB2001 against M^pro^ resistance variants. Clear examples of single amino acid substitution mutations conferring selective resistance to nirmatrelvir and ensitrelvir are highlighted in gray; similar mutations have yet to be found for FB2001. The relative values in brackets are reflected in the heatmap in Figure 1B.

Several single amino acid substitution variants including T21I, L50F, P252L, and T304I show minimal resistance to nirmatrelvir, ensitrelvir, or FB2001. Selective resistance to ensitrelvir is conferred by M49I and M49L, whereas selective resistance to nirmatrelvir is caused by A173V (highlighted in gray in **Table 1**). ΔP168 elicits similar resistance to all inhibitors, and synergistic resistance to nirmatrelvir when combined with A173V. S144A and L167F show the greatest resistance to ensitrelvir, intermediate resistance to nirmatrelvir, and lower resistance toward FB2001. In contrast to E166A and L50F/E166A, which cause a similar broad-spectrum resistance, E166V and L50F/E166V elicit very high resistance to nirmatrelvir, intermediate resistance to ensitrelvir, and substantially lower resistance to FB2001.

In addition to providing a method to rapidly profile candidate resistance mutations in living cells, our gain-of-signal assay also provides a quantitative metric for M^pro^ functionality^12^ (**Methods**). This system is based on the fact that overexpression of wildtype SARS-CoV-2 M^pro^ results in the cleavage of multiple substrates in cells^13,14^ including at least one required for RNA Polymerase II-dependent gene expression^12^. Therefore, expression of the Src-M^pro^-Tat-Luc reporter itself is rapidly shut down following transfection and can only be recovered by chemical or genetic inhibition of M^pro^. Thus, genetic mutations effectively phenocopy the chemical dose-responsiveness of the system, with some variations showing wildtype M^pro^ activity (background luminescence) and others compromising activity weakly or strongly depending on the nature of the mutation (low to high luminescence). For example, in comparison to wildtype M^pro^, catalytic mutants such as C145A yields 50- to 100-fold higher luminescence^2,12^. The M^pro^ variant constructs used here display a range of luminescence levels in the absence of drug indicative of near-normal M^pro^ activity (notably, M49I and M49L), weakly compromised M^pro^ activity (notably, A173V), and strongly compromised M^pro^ activity (notably, E166V) (**Figure S2**). These results suggest that several variants can confer at least partial drug resistance with little loss in M^pro^ functionality (and accordingly high viral fitness), whereas others such as E166V require suppressor mutations such as L50F to restore M^pro^ function to a level that enables virus replication (evidenced by recent resistance studies with pathogenic SARS-CoV-2 in culture and *in vivo* in animal models^3,5^).

Regardless of the details of each molecular mechanism, the results here demonstrate that nirmatrelvir, ensitrelvir, and FB2001 have distinct resistance profiles and that the latter inhibitors (with appropriate formulations) may be effective in patients suffering from Paxlovid rebound^15^ or *bona fide* resistance^2^. FB2001 may additionally have a higher resistance barrier given that no fully functional single M^pro^ variants tested to-date confer a strong resistance to this compound. Importantly, the gain-of-signal live cell assay recapitulates recent findings using replication competent viruses and provides a safe and rapid method for assessing resistance. As the SARS-CoV-2 variant pool deepens, this assay and variant panel can be expanded in lock-step to provide early resistance “fingerprints” of candidate next-generation M^pro^ inhibitors. Such an early profiling strategy has the potential to minimize the risks of developing drugs prone to cross-resistance and, importantly, to help identify inhibitors with the highest barriers to resistance.

## Acknowledgments

This work was supported by National Institute of Allergy and Infectious Disease grant U19-AI171954. RSH is an Investigator of the Howard Hughes Medical Institute, a CPRIT Scholar, and the Ewing Halsell President’s Council Distinguished Chair at University of Texas Health San Antonio.

