## Supplementary Methods and Figures S1-S2 for "Rapid resistance profiling of SARS-CoV-2 protease inhibitors"

Contents: Methods and Data Availability, Figures S1 and S2

### Methods and Data Availability

#### Cell culture

All M$^{pro}$ inhibition assays were done as described with the live cell gain-of-signal assay using the pcDNA5/TO-Src-M$^{pro}$-Tat-fLuc reporter construct[12]. All M$^{pro}$ single and double mutants selected for analysis here were based on recent reports of candidate resistant mutatnts[2-9] generated by site-directed mutagenesis (primers available upon request) and verified by Sanger sequencing. Transfections were done using 293T cells maintained at 37°C and 5% $CO_2$ in DMEM (Gibco catalog number 11875093) supplemented with 10% fetal bovine serum (ThermoFisher catalog number 11965084) and penicillin-streptomycin (Gibco catalog number 15140122).

**$M^{pro}$ resistance experiments**

For each individual $M^{pro}$ variant, $3x10^6$ 293T cells were plated in a 10cm dish and transfected 24h later with 2µg of the corresponding variant plasmid using TransIT-LT1(Mirus catalog number MIR 2304). Transfected cells were incubated at 37°C and 5% $CO_2$ for 4h, washed once with phosphate buffered saline (PBS), trypsinized, resuspended in fresh media, and diluted to a concentration of $4x10^5$ cells/ml. 50µL of each cell suspension was added to a 96-well white clear bottom cell culture plate (ThermoFisher #165306) containing pre-aliquoted inhibitor-supplemented media for a final concentration of 20,000 cells per well and inhibitor dose response range of 10µM to 2.4nM. Inhibitors were purchased from commercial vendors (nirmatrelvir, MedChemExpress catalog number HY-138687; ensitrelvir, MedChemExpress catalog number HY-143216; FB2001, Sigma-Aldrich catalog number SML2877) and purity was confirmed by HPLC and NMR. After an additional 44h incubation (48h total post-transfection), luciferase activity was quantified by removing growth medium and adding 50µL of Bright-Glo reagent (Promega catalog number E2610) to each well and incubating at room temperature in the dark for 2m before measuring luminescence on a Biotek Synergy H1 plate reader.

Percent $M^{pro}$ inhibition was calculated at each concentration of inhibitor using the formula below using the relative luminescence of an inhibitor (RLi) treated sample to the untreated control for each individual mutant.

$$\%\, inhibition = \%100 - (100/(RLi)$$

Results were plotted using GraphPad Prism 9 and fit using a four-parameter non-linear regression to calculate $IC_{50}$ values (**Figure S1**; **Table 1**). Resistance of mutants was calculated by the fold change in $IC_{50}$ of the mutant relative to WT $M^{pro}$, and these values were used to generate a heatmap in GraphPad Prism9 (**Figure 1B**).

As an increase in luminescence in the absence of any inhibitor treatment is indicative of decreased $M^{pro}$ catalytic activity, the relative activity of each mutant was calculated by the formula below using the relative luminescence of a mutant (RLm) to the WT enzyme in the absence of inhibitor (**Figure S2**).

$$\%\ activity = \%100 - [100/RLm]$$

**Data Availability**

All results are presented in the main display items or supplementary figures. The $M^{pro}$ gain-of-signal system is available upon email request to and completion of an appropriate MTA (U.S. Provisional Application Serial No. 63/108,611, filed on November 2, 2020).

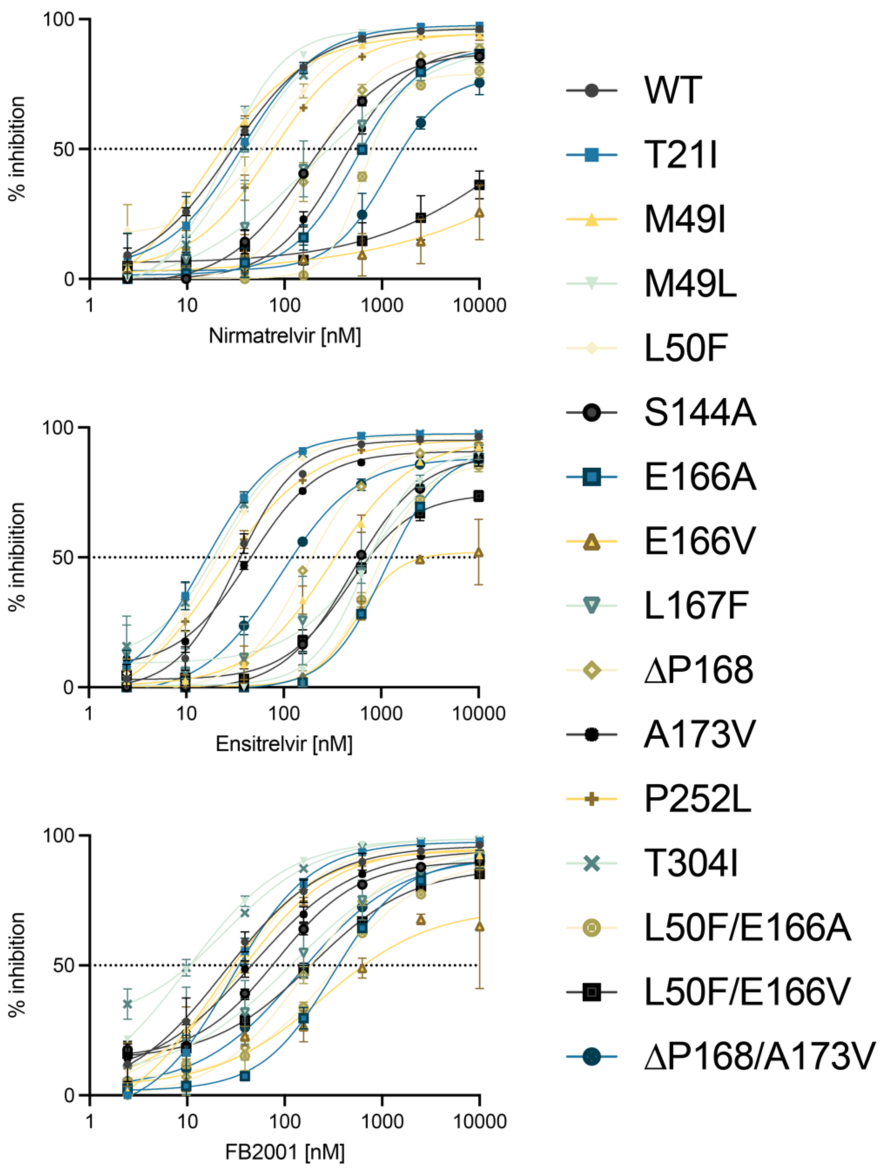


**Figure S1. Dose response curves showing inhibition of WT and mutant M$^{pro}$ enzymes by nirmatrelvir, ensitrelvir, and FB2001.** Dose response of respective M$^{pro}$ variants using the gain-of-signal assay in cells treated with indicated inhibitors in a 4-fold serial dilution beginning at 10µM (data are mean +/- SD of biologically independent triplicate experiments). $IC_{50}$ values for each inhibitor are listed in **Table 1**.

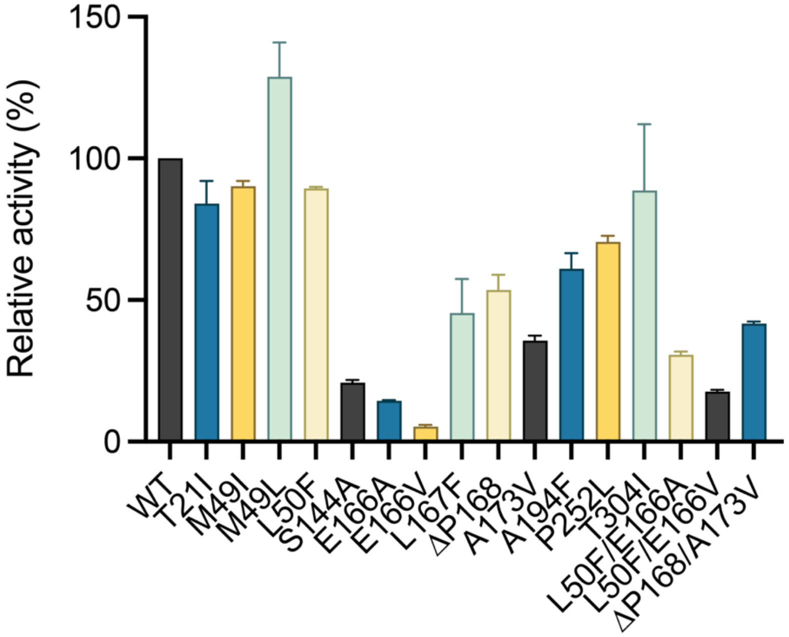


**Figure S2. Relative activity of $M^{pro}$ mutants.** A histogram showing the relative catalytic activity of each $M^{pro}$ mutant relative to the WT construct (normalized to 100% to facilitate comparison). Several single mutants such as T21I, M49I, M49L, L50F, and T304I show near WT activity. Other mutants such as A173V show modest 1.5 to 3-fold decreases in relative activity, and a few such as E166V are severely compromised. L50F partly restores the activity of E166A and E166V mutants consistent with prior reports[2-9].
